# General-Purpose Large Language Models, such as DeepSeek V3.2, Have Evolved Protein Design Capabilities

**DOI:** 10.1101/2025.11.23.689994

**Authors:** Jiawei Li, Xinxiu Dong

## Abstract

General-Purpose Large Language Models (GLLMs), although primarily developed for natural language processing, are increasingly demonstrating emergent capabilities in specialized scientific domains. In this study, we explored the potential of GLLMs,specifically DeepSeek V3.2 Exp in reasoning mode to perform practical protein engineering tasks without domain-specific biological training. Two representative design problems were addressed: Generation of amino acid sequences predicted to adopt the canonical 4- helix bundle topology, and targeted mutation design to improve protein solubility while preserving core structural integrity. Across 49 generated 4- helix bundle candidates, 40 adopted the desired geometry, with 36 achieving pLDDT scores above 70. Solubility optimization on 50 representative proteins yielded 46 mutants with an average predicted score increase of 0.178, and 29 maintained structural deviations below 3 Å RMSD. These results indicate that general-purpose LLMs such as DeepSeek V3.2 can integrate sequence–structure–property relationships sufficiently to produce viable protein designs. We propose a hybrid workflow that couples GLLM-based mutation generation with established computational validation, offering an accessible route for protein and peptide engineering.

## Introduction

Artificial Intelligence (AI) as a 4.0 industrial revolution is increasingly bringing smart solutions to humans^1^. In recent years, AI technologies—particularly general-purpose Large Language Models (GLLMs) such as ChatGPT, Claude, DeepSeek, Qwen, and Gemini—have experienced explosive growth. These models have been widely adopted by individuals and enterprises, demonstrating extensive applications in text generation, code development, customer service, document processing, and numerous other fields, thereby transforming workflows and fostering new forms of innovation.

In recent years, AI models tailored for protein systems have delivered remarkable breakthroughs, significantly advancing our ability to predict structures, design novel proteins, and understand complex biological functions.

Examples include AlphaFold 2^2^, which achieves near-experimental accuracy in predicting protein structures from sequence alone; ProteinMPNN^3^ and RFDiffusion^4^, which enable inverse folding and backbone generation for de novo design at unprecedented success rates; the ESM^5^ family of protein language models, which capture evolutionary and functional signals from hundreds of millions of sequences to support zero-shot mutation effect prediction and conditional protein generation; and MaSIF^6^ or PSICHIC^7^, which harness geometric and physicochemical deep learning for rapid binding-site identification and virtual screening.

Specialized models for protein have demonstrated remarkable performance in recent years, yet their accessibility remains limited in certain practical research contexts. For many wet-lab researchers, a lack of background in computational biology and programming skills — and even unfamiliarity with the Linux environment — can pose substantial technical barriers. Moreover, hardware resources, particularly high-performance GPUs, are not only scarce but also prohibitively costly. These factors make it difficult for such researchers to directly deploy and run the latest deep learning tools, thereby preventing them from fully benefiting from advances in artificial intelligence within the field of biology. Notably, this issue is being gradually alleviated through various approaches, such as cloud-based platforms like AlphaFold Server, which offer more convenient remote access. Furthermore, some emerging models, such as Pinal^8^, are beginning to explore natural language–driven interaction modes, thereby further lowering the barriers to use.

A peptide design competition permitting unrestricted methodologies and allowing a maximum of ten sequences per participant was undertaken. Due to incomplete design work prior to the submission deadline, the remaining five sequences were generated using DeepSeek v3.1 shortly before the cutoff on September 30, 2025. Notably, one of these sequences successfully passed AlphaFold3’s preliminary screening for competitiveness and inter-protein Template Modeling score (ipTM), and progressed to wet-lab validation, ultimately ranking 13th among 149 entries that advanced to the wet lab stage. This unexpected outcome prompted a preliminary investigation into the feasibility of employing GLLMs, without domain-specific biological training, for practical protein engineering tasks, and exploring prompt usage techniques.

One study case involved generating sequences predicted to fold into the 4-helix bundle architecture, a canonical structural motif composed of four closely packed parallel or antiparallel α-helices organized around a hydrophobic core and stabilized by hydrogen bonding and electrostatic interactions. This topology is widely present in natural proteins, including membrane transporters, signaling mediators, and certain enzyme domains. It is also a classic model for studying sequence–structure relationships^9^.

Another case addressed the optimization of protein solubility, a physicochemical property crucial for efficient expression, purification, and downstream applications such as drug development, industrial biocatalysis, and structural biology. Insufficient solubility often results in aggregation or precipitation, thereby impeding functional analysis and application. Introducing targeted mutations to enhance solubility is a common strategy in protein engineering and synthetic biology.

In addition, based on our practical experience during the experiments, we discussed the applicable scenarios and prompt usage techniques for employing GLLMs in protein sequence design, and established an online post dedicated to sharing prompts and design cases to facilitate communication, reproducibility, and collaborative improvement within the research community. Overall, the primary objective of this work is to conduct an early exploration into the feasibility of applying such models to protein design tasks, aiming to raise awareness of their practical availability, investigate effective usage strategies, and lower the technical barrier for researchers without computational biology expertise. For experienced practitioners, this approach may also offer a simpler and more efficient alternative for certain small-scale engineering problem.

## Method

IP address restrictions prevented access to certain large language model services, such as ChatGPT. Consequently, we selected DeepSeek, one of the most capable models previously used in our experiments and accessible under these network constraints. However, the official API for DeepSeek- V3.1 is no longer available, and therefore the present study adopted DeepSeek- V3.2- Exp — the highest-performance variant currently accessible via the official API.

All dialogues in this study were conducted using DeepSeek- V3.2- Exp in reasoning mode, accessed via the official DeepSeek API (https://platform.deepseek.com/) through a Python script. All requests used fixed parameters: the model was set to deepseek-reasoner (corresponding to DeepSeek- V3.2- Exp reasoning mode), with a message structure consisting of two components: A system prompt defining the scope and constraints of the task, and a user prompt specifying the details of the experimental task. Across all experiments, generation parameters were kept constant: temperature = 1, top_p = 1, presence_penalty = 0, frequency_penalty = 0, stream = False. Unless otherwise noted, prompts were written in English, in accordance with its widespread use in scientific communication and literature. Notably, DeepSeek has been specifically optimized for both English and Chinese, allowing efficient handling of prompts and outputs in either language.

All figures were generated using Matplotlib^10^ (v2.2.3), with a custom gradient color map ranging from orange to blue to represent predicted Local Distance Difference Test (pLDDT) values.

### Generation of 4-helix bundle structures

For this target fold, standardized system and user prompts (see Supplementary Materials) were prepared to instruct the model to generate amino acid sequences predicted to adopt the 4-helix bundle geometry. For each target structure, a single sequence was generated per model call, and the task was independently repeated 50 times to achieve a statistically meaningful dataset. All sequences were submitted to ColabFold^11^ (AlphaFold2^2^) for structure prediction with default parameters. Structural evaluation consisted of visual inspection of secondary structure organization and spatial helix arrangement using PyMOL, to confirm conformity to the bundle motif; and computation of the pLDDT score to quantify confidence in the predicted structures.

### Protein solubility optimization

In this experiment, 50 representative proteins were randomly selected from the MTPSol^12^ dataset as initial inputs. The model was instructed to propose amino acid mutations aimed at increasing solubility while retaining the original core fold. Generated mutant sequences were evaluated using the CamSol^13^ (https://www-cohsoftware.ch.cam.ac.uk/index.php/camsolintrinsic) algorithm to calculate their predicted solubility scores, which were compared with those of the wild-type sequences to assess improvement. Subsequently, the AlphaFold Server (based on AlphaFold3^14^) was used with default parameters to predict the structures of both mutant and wild-type proteins.

The Root Mean Square Deviation (RMSD) between mutant and original structures was calculated to quantify structural alterations. RMSD computation was performed by aligning the predicted structures using backbone Cα atoms as reference coordinates. Structural alignment and RMSD calculation were performed using the Superimposer module of BioPython^15^ (v1.81).

### Data

All datasets used in this study, all data generated, and all related code will be stored in the following GitHub repository: https://github.com/LIJIAWEI040301/GLLMs_for_protein

## Results

### Generation of 4-helix bundle structures

A total of 49 sequences were successfully generated, and their structures were predicted using ColabFold. Notably, 40 of these sequences had pLDDT scores above 70. Among all 49 generated structures, 40 successfully adopted the 4-helix bundle structure, with 36 of these structures having pLDDT scores greater than 70 (see Fig. 1). These results indicate that the model is highly reliable in predicting 4-helix bundle structures.

**Figure 1.**
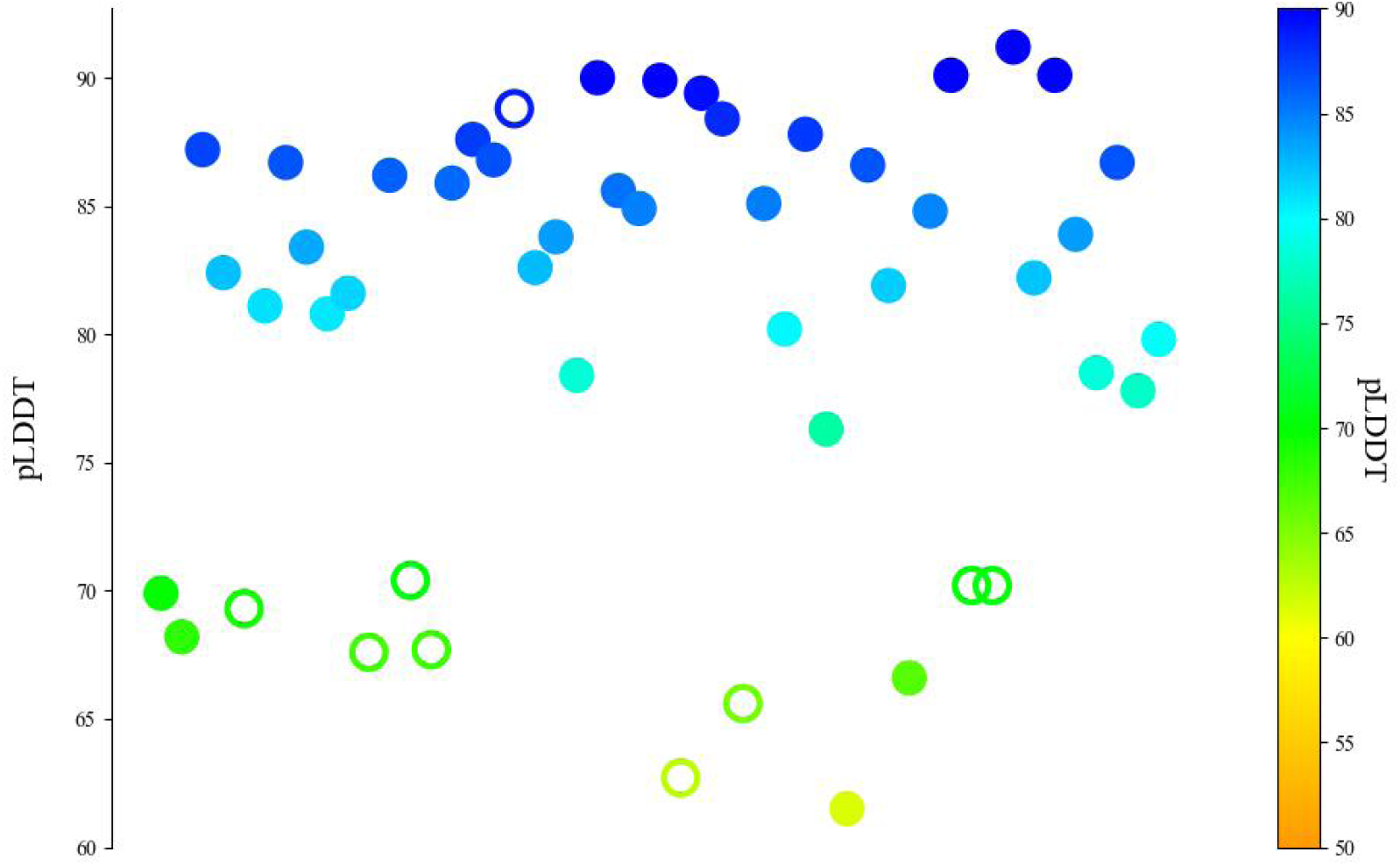
Generation outcomes of GLLM-designed 4- helix bundle proteins. Predicted pLDDT scores for 49 protein sequences generated by DeepSeek V3.2 Exp targeting the 4- helix bundle topology. Filled circles indicate sequences that successfully folded into the intended 4- helix bundle architecture based on AlphaFold2/ColabFold predictions and visual inspection, whereas open circles represent sequences that failed to adopt the target fold. A pLDDT score above 70 generally corresponds to high structural confidence.

### Protein solubility optimization

Among the 50 representative proteins examined, the model successfully generated 46 mutant sequences that met the design requirements. According to predictions from the CamSol algorithm, all of these mutants exhibited improved solubility scores compared with their wild-type counterparts, with an average increase of 0.178. Structural comparison revealed that 29 mutants showed relatively small conformational changes (RMSD < 3 Å), and 17 of them exhibited even smaller deviations (RMSD < 1 Å) (see Fig. 2). These results indicate that, even in the absence of domain-specific training in protein science, a general-purpose large language model can, in most cases, successfully generate mutant proteins with optimized physicochemical properties and high structural fidelity, but the approach is not stable.

**Figure 2.**
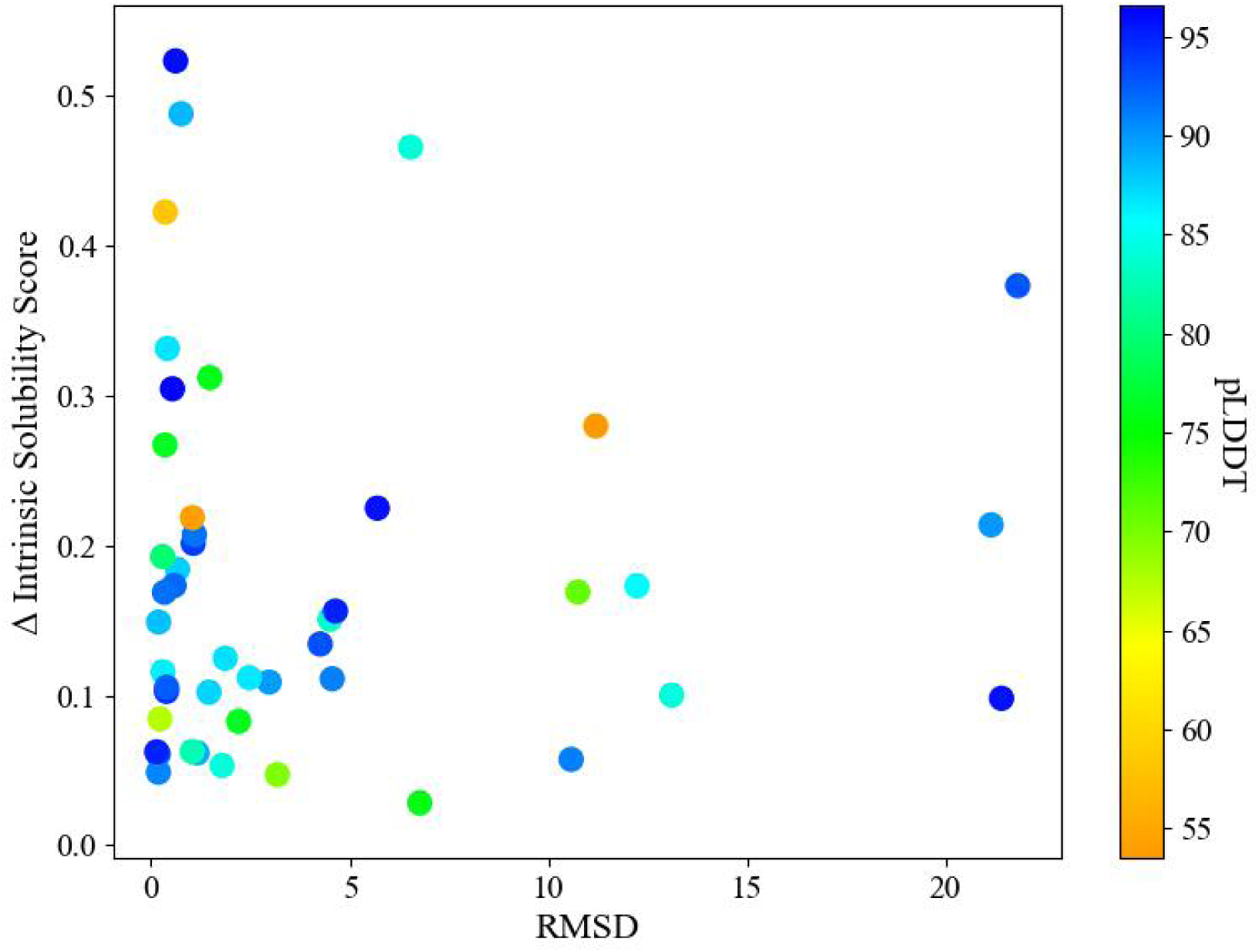
Solubility optimization and structural deviation of GLLM-generated protein mutants.

## Discussion

This study utilized the general-purpose large language model DeepSeek V3.2 Exp to successfully accomplish two protein engineering tasks: 4-helix bundle structure generation and protein solubility optimization. These results indicate that, even without specialized training in protein science, general-purpose large language models (such as DeepSeek) have already evolved certain protein engineering capabilities, enabling them to achieve meaningful outcomes in areas such as structural generation and physicochemical property optimization.

(Based on an analysis of the model’s reasoning content provided in the supplementary materials) General-purpose large language models may derive this capability from the large-scale, cross-domain knowledge acquired during training. Although the training corpus was not specifically tailored for protein science, it likely contained substantial amounts of publicly available content—such as molecular biology, chemistry, and structural biology resources, including academic publications and database entries—which enabled the model to implicitly learn associations between amino acid sequences, structural motifs, and physicochemical properties. Furthermore, the long-form reasoning capability of DeepSeek V3.2 Exp in reasoning mode, combined with its distinctive architectural design and optimization strategies, greatly enhances its ability to leverage and integrate such cross-domain knowledge. This allows the model, during protein sequence generation, to more effectively satisfy both structural topology requirements and physicochemical property optimization objectives, thereby demonstrating considerable potential for protein engineering applications. At the same time, the reasoning text generated by the model during the design process constitutes highly valuable interpretable data. Analyzing these reasoning contents can help researchers understand the model’s design rationale and, to some extent, assess the logical quality and plausibility of the generated sequences.

It also exhibits certain shortcomings similar to those observed when using general-purpose large language models to generate general textual information. For example, in the 4-helix bundle structure generation task, the model occasionally falls into repetitive patterns, continuously outputting the same tokens, as illustrated by the “design_040_design.txt” case in the supplementary file. Although such issues can often be easily resolved by re-running the generation process, they nonetheless occur. In addition, general-purpose large language models show instability when handling tasks that require precise positional annotations—their weakness in “counting” still manifests in protein engineering contexts. For instance, in the solubility optimization task, the model outputs mutation instructions in formats such as “A38E,” but occasionally produces incorrect residue numbering or omits mutation information, leading to failures in the design workflow. These observations suggest that, although the model does indeed possess protein engineering capabilities, its reliability and stability remain current weak points.

Similar to other types of GLLMs applications, prompt design strategies have a significant impact on model performance. According to the official DeepSeek prompt usage guidelines^16^ and our own usage experience in this study, evaluation results indicate that the model is highly sensitive to prompt content. In most cases, few-shot prompting is less stable than zero-shot settings. The zero-shot approach—directly describing the task objectives and specifying the output format—usually yields the best performance. Unless preliminary testing or reasoning analysis confirms that the model indeed lacks the necessary domain knowledge, it is not recommended to include redundant background information or external knowledge in the prompt.

We further found that the “reasoning length per amino acid residue”—that is, the total generated sequence length divided by the model’s internal reasoning steps—has a decisive influence on generation quality. This is easy to understand: reasoning resources are limited, and the model tends to compress its thought process. The longer the sequence, the less reasoning budget can be allocated per residue under the same constraints, leading to a decline in output quality.

In terms of prompting methods, directly asking the model to generate a complete protein sequence often results in missing outputs, unexpected mutations, or structures that do not meet design objectives. This not only affects sequence accuracy but also reduces the per-residue reasoning budget. In early mutation experiments, we adopted the full-sequence output strategy, and approximately half of the sequences exhibited these undesirable issues. In contrast, converting the task into position-specific mutation instructions (e.g., “A38E” format) significantly reduced the error rate and greatly improved result stability.

When directly replacing part of traditional computational biology tools, although GLLM performance is comparable, stability is noticeably inferior; therefore, it is not the best choice unless necessary. Even when used, multiple runs and expert judgment are required to assess the results. However, in certain niche tasks, GLLMs demonstrate distinct advantages—particularly in expanding short peptide mutation libraries. For example, for a peptide of length 15, the single-site saturation mutation space can still be exhaustively enumerated, but when multiple mutation sites or combinations are involved, the theoretical search space expands rapidly (in some cases reaching tens of thousands of possibilities). In such scenarios, GLLMs can propose targeted mutation suggestions based on physicochemical properties, followed by screening with established computational validation methods (such as AlphaFold structure prediction and molecular dynamics simulations), thereby forming an efficient and practical protein/peptide design pipeline. This task inherently involves short sequences (allowing more reasoning resources per residue), and the integration of mature computational validation methods can partially offset the stability disadvantage while fully leveraging the model’s strength in efficient mutation generation. This is the effective pathway we summarized from the case described in the Introduction.

This direction has the potential to develop into a method with significant practical application value and relatively straightforward implementation. To facilitate the exchange, validation, and continuous optimization of related approaches, we have established a Reddit community at https://www.reddit.com/r/GLLMs_for_protein/ dedicated to sharing prompt design templates and representative case analyses. The community aims to assist researchers in exploring and applying this methodology more efficiently.

### Limitations

This study has several limitations. First, due to network and platform constraints, the range of available models was restricted, and a systematic comparative analysis among different types of GLLMs was not conducted. Second, the evaluation of structural and physicochemical properties relied heavily on computational prediction tools such as AlphaFold and CamSol, and their results may deviate from actual experimental outcomes. In addition, the tasks explored were relatively limited in scope and did not encompass more complex objectives.

### Future Work

Future research will focus on optimizing prompt design strategies, systematically exploring more effective usage methods, and continuously maintaining and enriching the established community platform to facilitate the sharing, validation, and refinement of related approaches.

## Conclusion

In this study, the general-purpose large language model DeepSeek V3.2 Exp was successfully applied to two representative protein engineering tasks: generation of the 4- helix bundle structure and optimization of protein solubility. The results demonstrate that general-purpose large language models possess certain capabilities for protein design and optimization. Through practical applications, we have summarized an efficient workflow for small-scale protein/peptide design, which lowers the technical barrier while providing feasible pathways and new ideas for exploring innovative design strategies. This workflow shows potential applicability and scalability in relevant fields.

